## Supplementray Information for "Interactions between culturable bacteria are predicted by individual species’ growth"

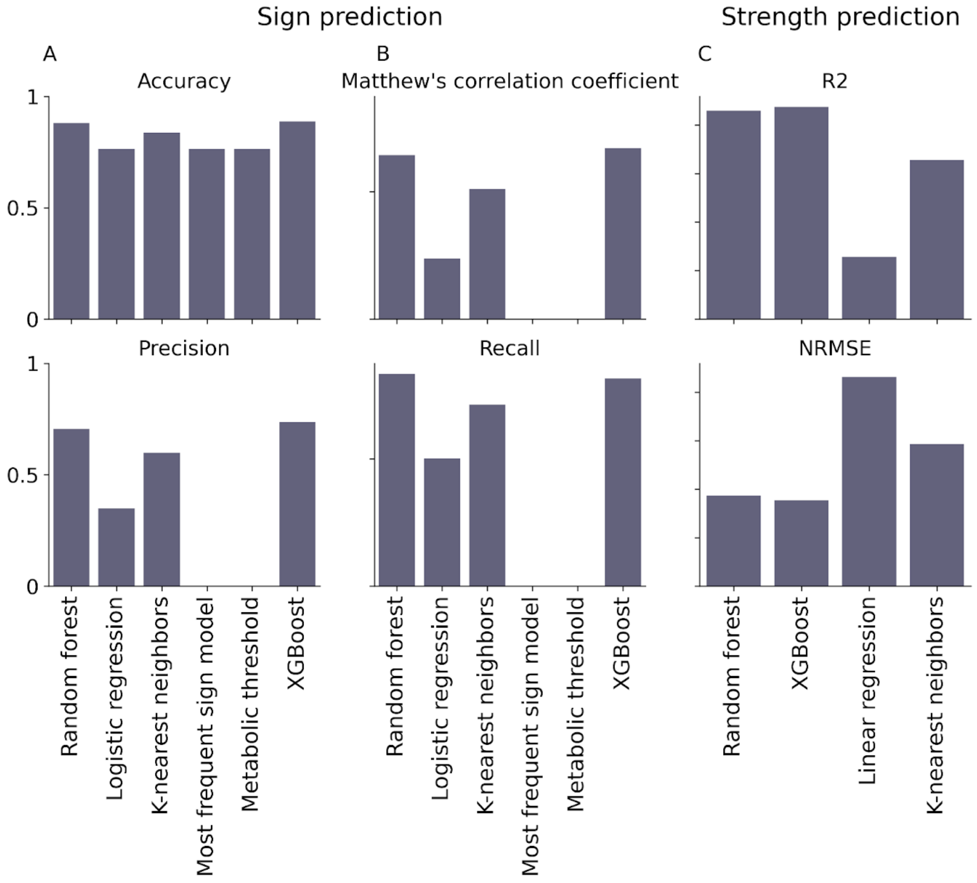


***S1. XGBoostclassifier outperformed all other evaluated models.*** *Comparison between all examined machine learning models for sign and strength predictions. Hyperparameters for each of the models (Random forest, XGBoost, linear regression, K-nearest neighbors) were tuned using RandomGridSearch (*scikit-learn 1.0.1*). For sign predictions, Accuracy, Matthew's correlation coefficient, precision and recall were compared. For strength predictions* $R^2$ *and NRMSE (normalized RMSE) were compared.*

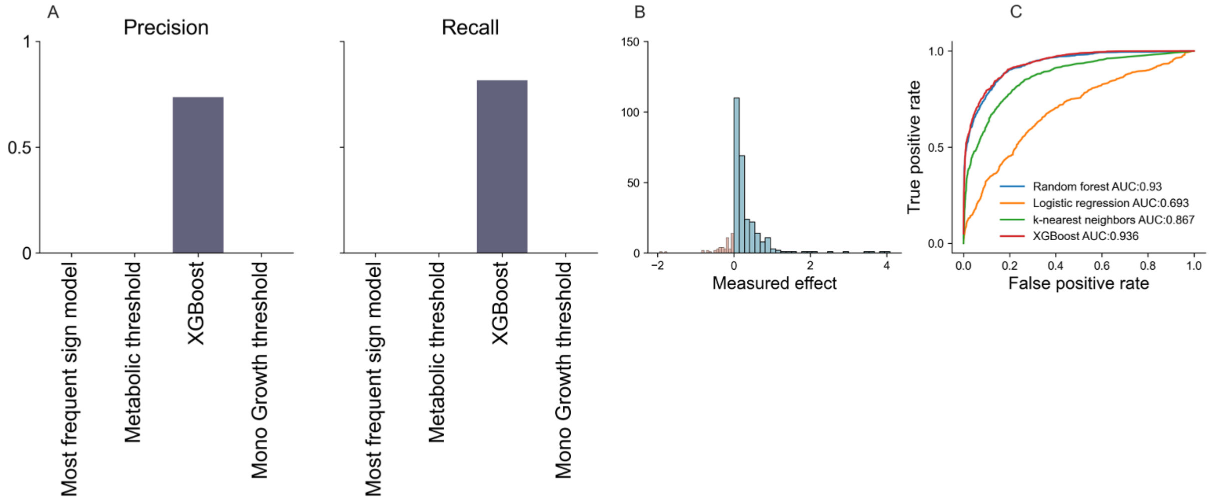


***S2. XGboost outperformed null sign prediction models. A.*** *Comparison of precision and recall across the best sign model null models (see Methods)* ***B.*** *Number of false positives(red) and false positives (blue) of the best sign prediction model as a function of the true effect strength.* ***C.*** *ROC curve of all machine learning models for effect sign predictions.*

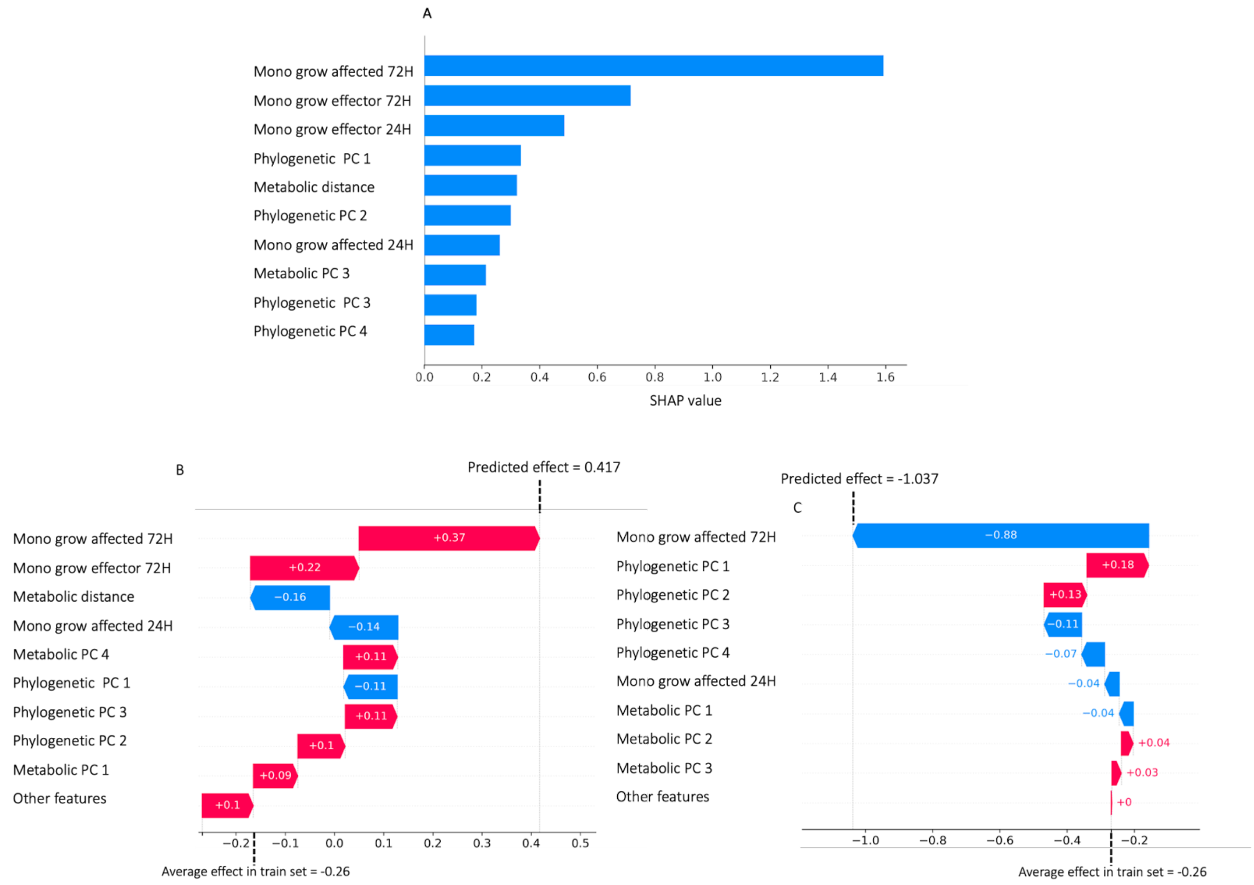


***S3. Monoculture growth yields of both species contribute the most to strength and sign predictions. A.*** *SHAP values for sign predictions. Samples labeled as positive (positive effect) are in blue, and samples labeled as negative (negative effect) are in red.* ***B.*** *Example of features' contribution to a single positive output prediction. X-axis shows the average effect in the train set (null strength prediction), and the y-axis shows the different features and their values (contribution) to the output of the model.* ***C****. Example of features' contribution to a single negative output prediction. X-axis shows the average effect in the train set (null strength prediction), and the y-axis shows the different features and their values (contribution) to the output of the model.*

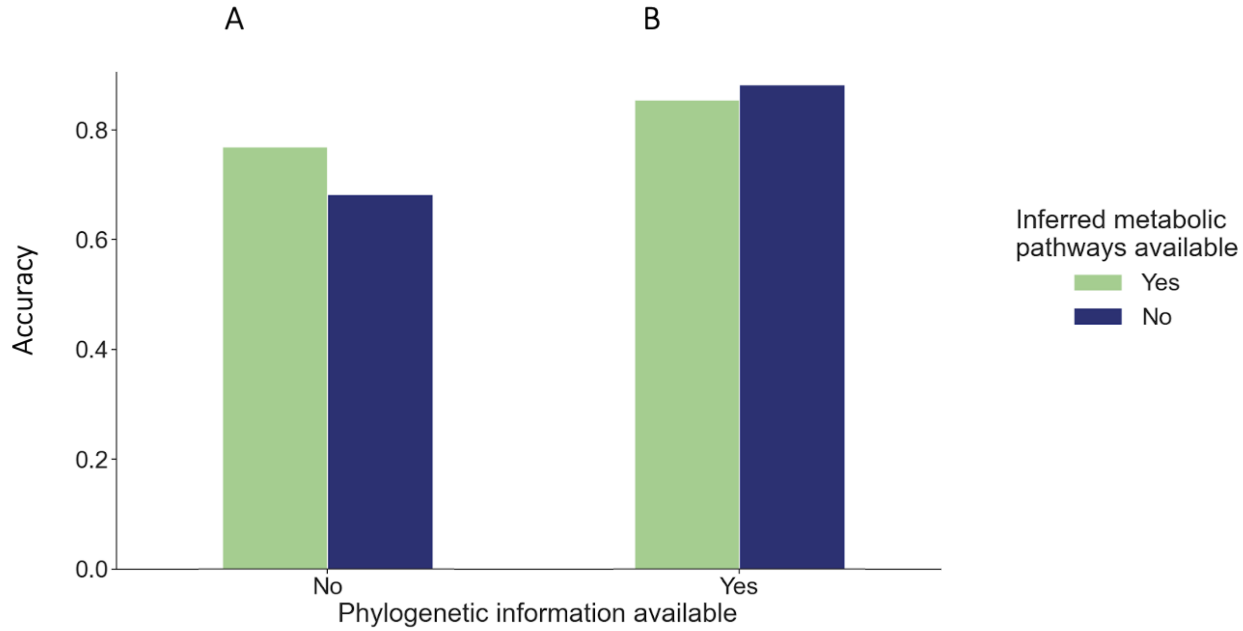


***S4.* When phylogenetic information was available, inferred *species' metabolic pathways did not improve predictions.*** *We tested whether information about species metabolic pathways can improve predictions of interaction sign. To do so, we followed the approach of DiMucci et al*[1]*. Briefly, we inferred the metabolic pathways encoded by each species based on 16S sequences using picrust2. Each metabolic pathway was encoded as a binary feature, indicating whether a metabolic pathway occurs in a species. In addition to information about the metabolic pathways, carbon sources were represented using binary vectors (one-hot encoding), together with features representing properties of the carbons sources, (number and weight of atoms ,*supplementary *Table 2). Lastly, species phylogeny was captured by the first two principal components of the phylogenetic distance matrix.*

***A.*** *Information about the inferred metabolic pathways increased prediction accuracy when the phylogenetic features were not included, consistent with previous findings*[1]*.* **B.** *However, when the phylogenetic features were included, a similar prediction accuracy was achieved with and without information about the inferred metabolic pathways.*

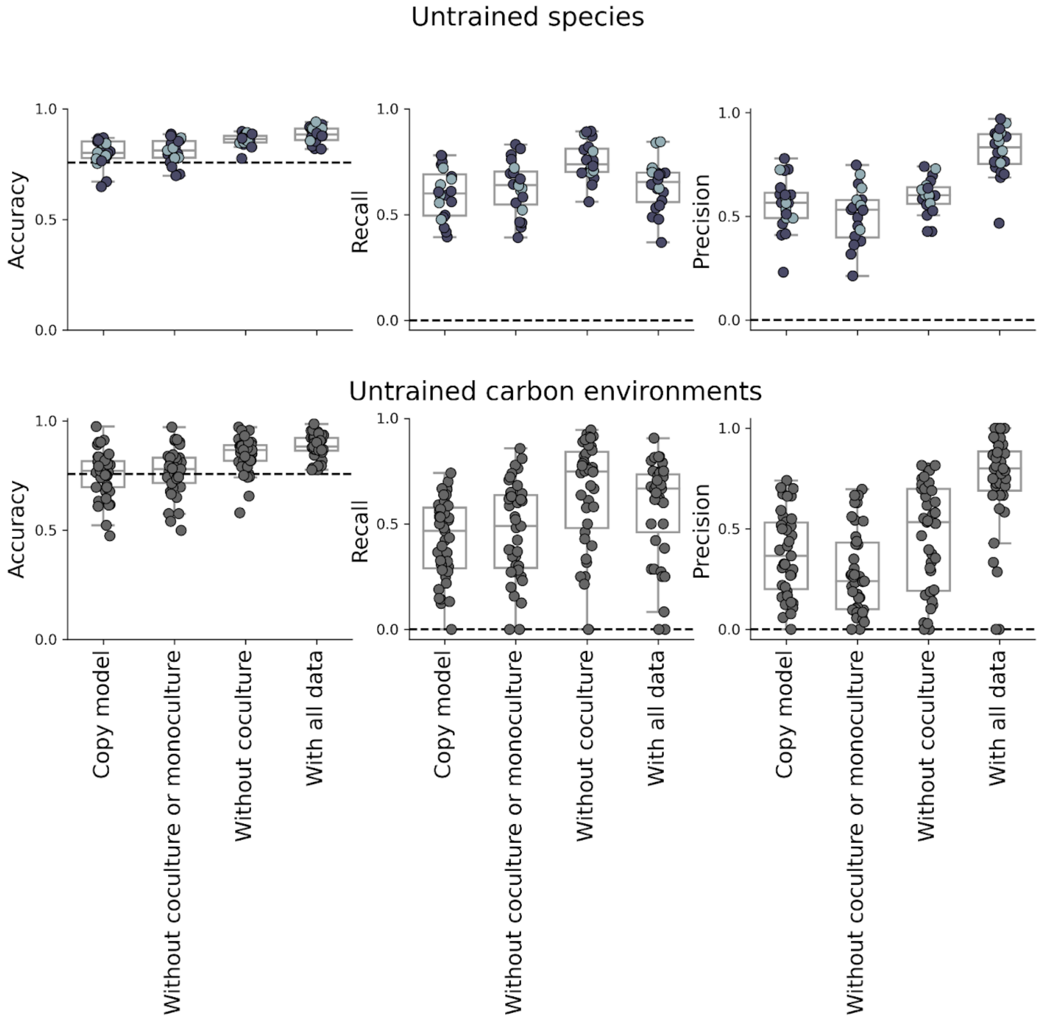


**S5. *Performance of models trained using partial information regarding a species or carbon environment.***

*The models differ by the features they are trained on and their test set (see Methods). Copy model can be either a phylogenetic copy model for "uncultured species", or a metabolic copy model for "uncultured" environments (see Methods). Other models are as described in Fig. 4. Dashed line is the score of a null sign model/null strength model.*

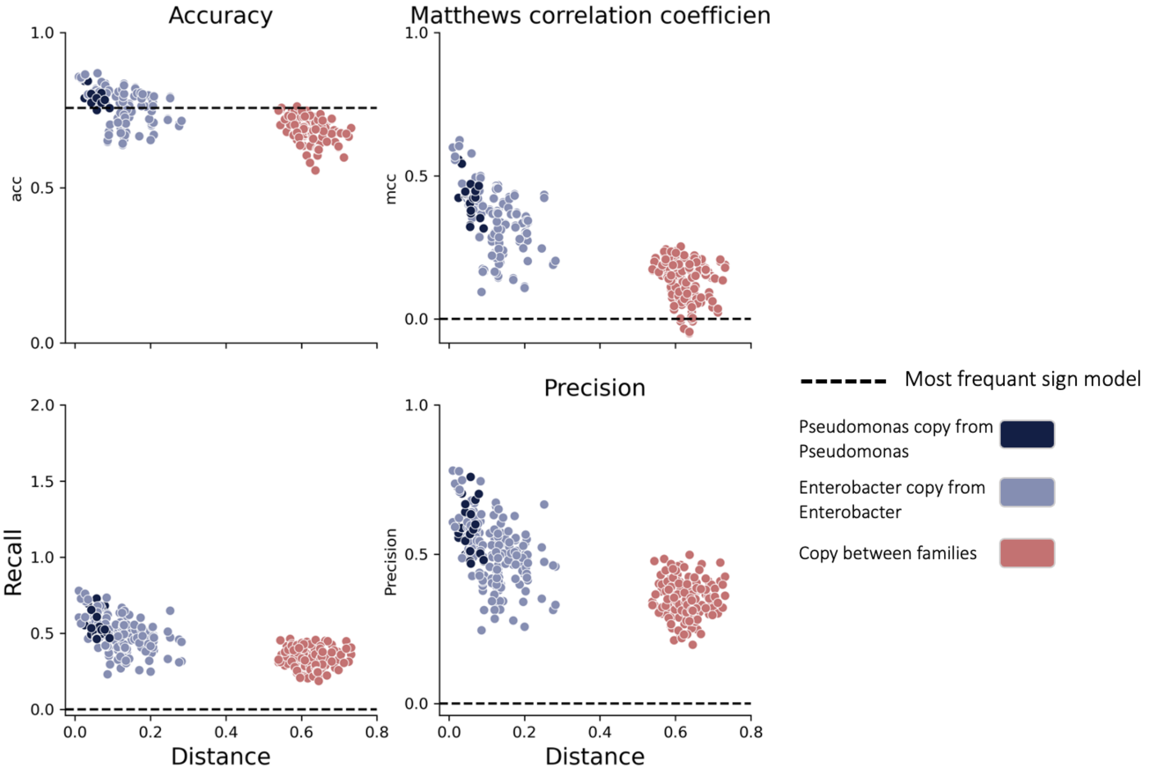


***S6. A simple phylogenetic copy model can provide predictive power when based on closely related species, in all examined matrices (Accuracy, Recall, MCC and precision).*** *Each dot represents a prediction for a single species, using one of the other 19 species as the copied species (interaction of the copied species in the same carbon source.) Different colors represent the distance category: the copied strain is from the same taxonomic family (dark blue, light blue) or form a different taxonomic family (red).*

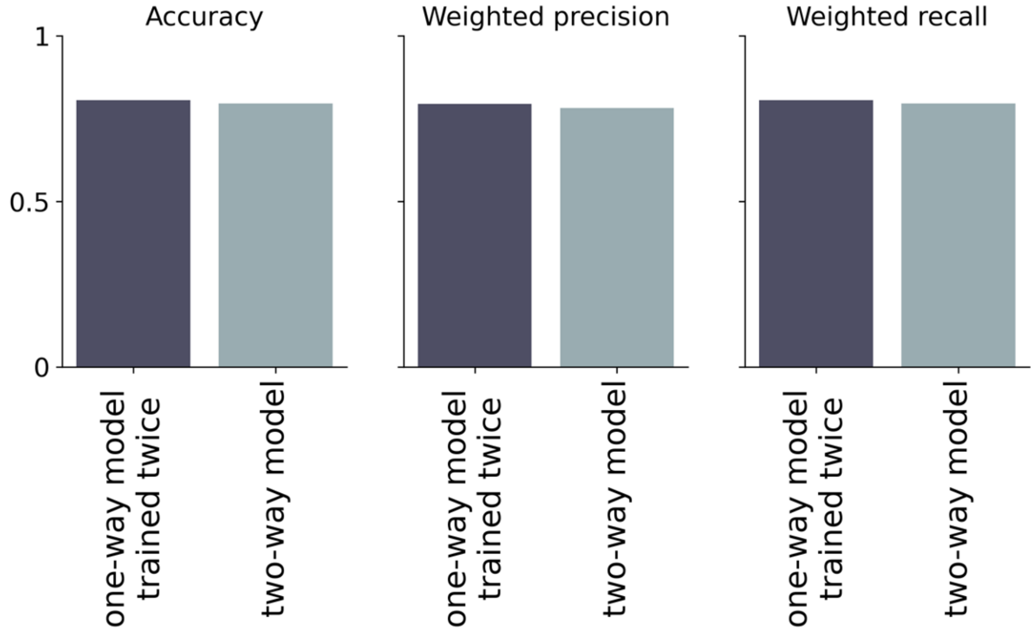


***S7*. *Jointly predicting reciprocal effects was not more accurate than combining two independent one-way effect predictions.*** *One-way models (both sign and strength) were trained twice, on both effects separately. Two-way model was trained on both effects at the same time (multilabel prediction). All trained models (both one-way and two-way) are with the same parameters* ***A.*** *Accuracy and weighted average of recall and precision for sign predictions of trained two-way (multilabel) model vs one-way model which is trained twice.*

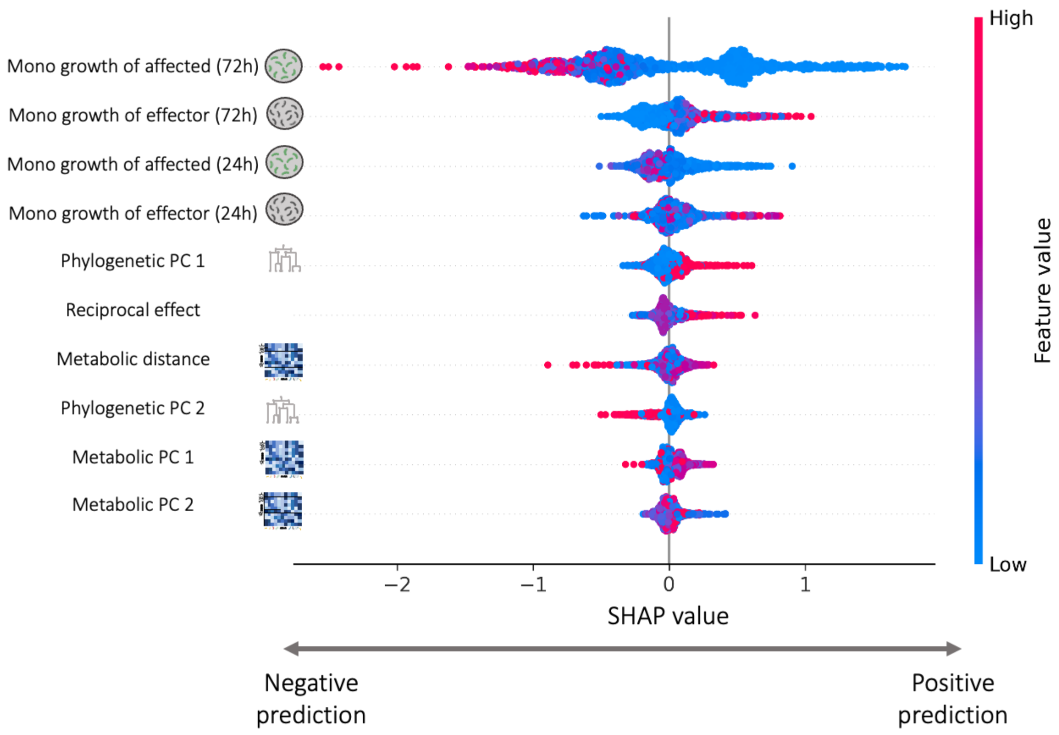


***S8. Top 10 most important features in one-way strength model with the reciprocal effect as a feature.*** *High SHAP values indicate positive influence on the predicted effect, and low values indicate negative influence on the predicted effect. Y-axis represents the different features, sorted in descending order according to their contribution to the model. Each dot represents a simulation of the model with a single change at a single feature value. The different colors represent the value of the feature.*

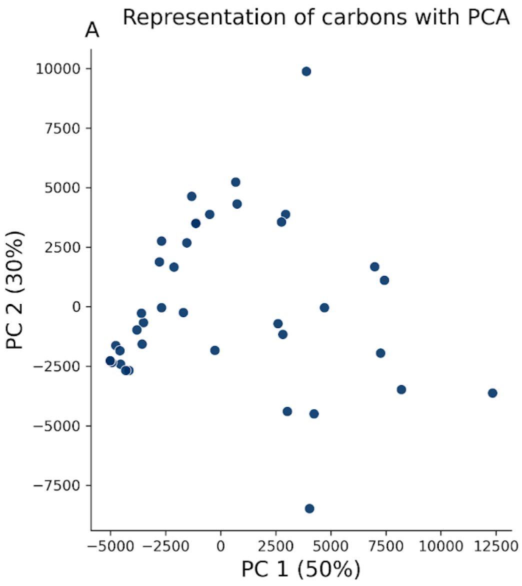


***S9. Carbon sources do not form distinct groups based on the species' growth profiles. A.*** *2D representation of carbon sources using the first two PCs. Each dot represents a single carbon source. PCA was performed based on the metabolic profiles (monoculture growth yield of different species) of all carbons.* ***B.*** *2D representation of species using the first two PCs. PCA was performed based on the species' phylogenetic distance matrix.*

1. DiMucci, D., Kon, M. & Segrè, D. Machine Learning Reveals Missing Edges and Putative Interaction Mechanisms in Microbial Ecosystem Networks. *mSystems* **3**, e00181-18 (2018).
